# Sublethal heat reduces overall reproductive investment and male allocation in a simultaneously hermaphroditic snail species

**DOI:** 10.1101/2023.03.15.532738

**Authors:** Shanna van Dijk, Valentina Zizzari, Joris M. Koene, Yumi Nakadera

## Abstract

A well-known effect of sub-lethal temperature exposure in a diversity of species is a decrease in reproductive performance. Although this effect has been particularly emphasized for males or male reproductive functioning, it remains to be firmly demonstrated that the effect of heat on fertility is sex-specific. To contribute to this question, here we examined the impact of sub-lethally high temperature on male and female functions in a simultaneously hermaphroditic snail species, *Lymnaea stagnalis*. Examining hermaphrodites is useful to evaluate the sex-specific impacts of heat exposure, since they possess male and female functions within a single individual, sharing genetic and environmental factors. Moreover, previously developed sex allocation theory allows us to compare the differential performance of sex functions. In this study, we exposed snails to temperatures ranging from 20 to 28 °C for 14 days and assessed their egg and sperm production, sperm transfer, mating behaviour and growth. Both types of gamete production were significantly reduced by higher temperature, leading to an overall reduction of reproductive investment. By quantifying sex allocation, we furthermore revealed that the heat-stressed snails reduced the relative investment in their male function. In addition, even though sperm production and its transfer were drastically decreased by high temperature, male mating motivation was not affected. This study illustrates that examining simultaneous hermaphrodites can provide significant insights for the impact of heat, and the proximate mechanism, on reproduction in wildlife.

## Introduction

Evaluating the effects of elevated temperature on wildlife is an urgent task. It has been well established that not only the temperature at which an organism dies, i.e., Critical Thermal Limit (CTL), but also that the temperature at which fertility is lost, i.e., Thermal Fertility Limit (TFL), are crucial to determine the long-term viability of natural populations that experience increasing environmental temperatures (reviewed in Walsh et al. 2019a). For instance, Parrat et al. (2021) demonstrated that, across 43 *Drosophila* species, TFLs are commonly lower than CTLs, and the estimates of TFL greatly improve the model prediction on species distribution, implying that we are underestimating the impacts of global warming on natural populations. In addition, the reduced fertility under influence of heat has been reported in diverse plants and animals (e.g., Oshino et al. 2007; Sage et al. 2015; Paxton et al. 2016; Zizzari and Ellers 2011; Hurley et al. 2018; Sales et al. 2018; Stürup et al. 2013) including humans (e.g., Yogev et al. 2004). To date, however, it remains controversial which sex is more vulnerable to increasing temperatures (Iossa 2019; Walsh et al. 2019b). Based on several hypotheses, such as descended testicles in mammals, male or male reproductive functions are often the primary target to estimate TFLs. However, females also show reduced fertility under sublethal heat exposure (e.g., Blanckenhorn et al. 2014; Paxton et al. 2016: Zizzari and Ellers 2014). Although the sex-specificity of reduced fertility under heat would significantly improve the understanding of the physiological mechanisms causing such reduction, as well as their evolutionary consequences in wildlife, the experimental insights are limited and need urgent expansion (Prasad et al. 2011; Paxton et al. 2016; Park et al. 2017).

Contrasting with separate-sexed species, simultaneous hermaphrodites (hereafter hermaphrodites) represent an optimal system to test which sex is more vulnerable against heat exposure. Since these organisms possess functional male and female reproductive systems within a single body, we can determine the effects of heat on both sex functions within the same individuals. Hence, in contrast to separate-sexed species, hermaphrodites allow us to compare the sex-specific effects of heat on reproduction without cofounding effects, such as the influence of differential hormonal and genetic components, or sex-limited expression (e.g., Pélissié et al. 2012). In addition to this logistical advantage, the well-established theoretical framework to study and quantify sex allocation in hermaphrodites - how reproductive resources are allocated to the male or female function (e.g., Charnov 1979, 1982; Schärer 2009) – offers predictive power. Obviously, the backbone of sex allocation theory is selection and adaptation – the theoretical framework aims to understand why organisms change sex allocation under certain social or environmental condition, as the consequence of selection over evolutionary time. However, sex allocation theory can equally well be applied to evaluate the impact of heat on reproduction in hermaphrodites, with the significant caveat that any observed changes in sex allocation under the influence of temperature are not necessarily adaptive.

To the best of our knowledge, however, there are barely any studies that examined the impact of heat on male and female functions in hermaphrodites, asking, for example, which sex is more vulnerable to heat, or if sex allocation changes depending on temperature. For example, in a self-fertilizing hermaphroditic fish, *Kryptolebias marmoratus*, Park et al. (2017) showed that fish exposed to high temperature had a relatively smaller gonadosomatic ratio, but not testis area. Further investigation revealed the disruption of hepatic vitellogenin synthesis at high temperature, which led them to conclude that high temperature affects ovarian development more than testicular development. In a sequentially hermaphroditic self-fertilizing nematode *Caenorhabditis briggsae*, the results indicated that the declined fertility under higher temperature is mostly due to the compromised fertility of developing sperm, but not oogenesis nor sperm count (Prasad et al. 2011). Lastly, in the hermaphroditic snail *Bulinus truncatus*, the exposure to high temperature during development tends to produce snails without a functional penis (Schrag and Read 1992). Even though such aphallic individuals produce sperm and self-fertilize, they cannot transfer ejaculate to partners. Despite the lack of male mating ability, aphallic snails were found to be as fecund as euphallic individuals and no difference in sex allocation was observed (e.g., Doums and Jarne 1996; Doums et al. 1998). Although the number of studies is fairly limited and all the study species were self-fertilizing hermaphrodites (indicating their male investment is low, regardless of temperature), previous studies indeed suggest that examining hermaphrodites could tell which sex function is more affected by heat, motivating us to examine the impacts of elevated temperature on hermaphrodites and their sex allocation.

Given the above, we examined the impacts of heat exposure on reproduction in a hermaphroditic species, *Lymnaea stagnalis*. It has been well established in this species that exposure to higher temperatures alters egg production as well as immune response (Leicht and Seppälä 2019; Salo et al. 2019; Leicht et al. 2013, 2017; Seppälä and Jokeka 2011), memory formation (Teskey et al. 2012; Sunada et al. 2016), food consumption (Zhang et al. 2018), synaptic transmission (Sidorov 2002), respiration (Sidorov 2012) and so forth (Salo et al. 2017). Also, the reproductive biology of *L. stagnalis* is well studied and their reproductive performances are readily quantifiable (e.g., sperm count: Loose and Koene 2008, fecundity: van Iersel et al. 2014, mating behaviour: Nakadera et al. 2015; Moussaoui et al. 2018). In this study, we focused on evaluating the difference between male and female functions, by comparing egg and sperm production by the same individuals. In addition, we measured the effect of heat on growth and mating behaviour. We report here that the snails exposed to 28 °C for two weeks reduced their overall investment in reproduction and their male function was diminished more than their female function. In addition, we revealed that their drastically reduced sperm production did not discourage them from mating in the male role.

## Material and Methods

We used adult *L. stagnalis* from the long-standing lab culture at Vrije Universiteit Amsterdam. During rearing, these snails were kept in flow-through tanks with aerated low-copper water at 20 ± 1 °C and a light:dark cycle of 12:12h. They were fed with broadleaf lettuce and fish flakes (Tetraphyll, TetraGmbH) *ad libitum*. We used an age-synchronized cohort of snails, which was three-month-old at the start of the experiment and fully sexually matured as evidence by their egg laying capability. This species is simultaneously hermaphroditic and shows unilateral mating, meaning that, in a single mating event, one snail acts as male (sperm donor) and its partner as female (sperm recipient). When motivated, they can swap their sex roles immediately after their first mating (Koene and Ter Maat 2005).

### Heat exposure setup

We randomly selected 48 snails to expose to 20 °C, 24 °C and 28 °C for 14 days, respectively (Fig. 1). We set 20 °C as control, because this is the standard rearing temperature and near their thermal optimum (Vaughn 1953; Van der Steen et al. 1969; Nakadera et al. 2015). We chose 24 °C and 28 °C as simulated warm and extremely warm summer conditions in shallow water bodies (see also Salo et al. 2019). We chose to examine the effect of continuous heating for 14 days, rather than a brief heat shock, as water is slow to increase or decrease its temperature compared to air. Moreover, this duration is long enough to evaluate the effect of heat on sperm production, since their spermatogenesis takes less than 10 days (De Jong-Brink et al. 1985, Nakadera et al. 2020). To achieve these heat treatments, we used six aquariums (ca. 15 L) with heaters. These aquariums have a slow flow-through of aerated water, similar to the standard breeding tanks. Within an aquarium, we placed 8 perforated containers (400 ml) to monitor the snails. To acclimate the snails to a designated temperature, we put snails in a closed container with water at 20 °C, and placed the container in the aquarium for a few hours until the water inside the container had the same temperature as the surrounding aquarium water. Then, we exchanged the closed containers with perforated containers to initiate heat exposure. Throughout the experiment, we fed the snails with a lettuce disc (ca. 19.6 cm^2^) per day per capita. Due to the limited number of available aquariums with individual thermostats, we ran the same experiment twice to ensure a high enough sample size (N = 96).

**Fig 1.**
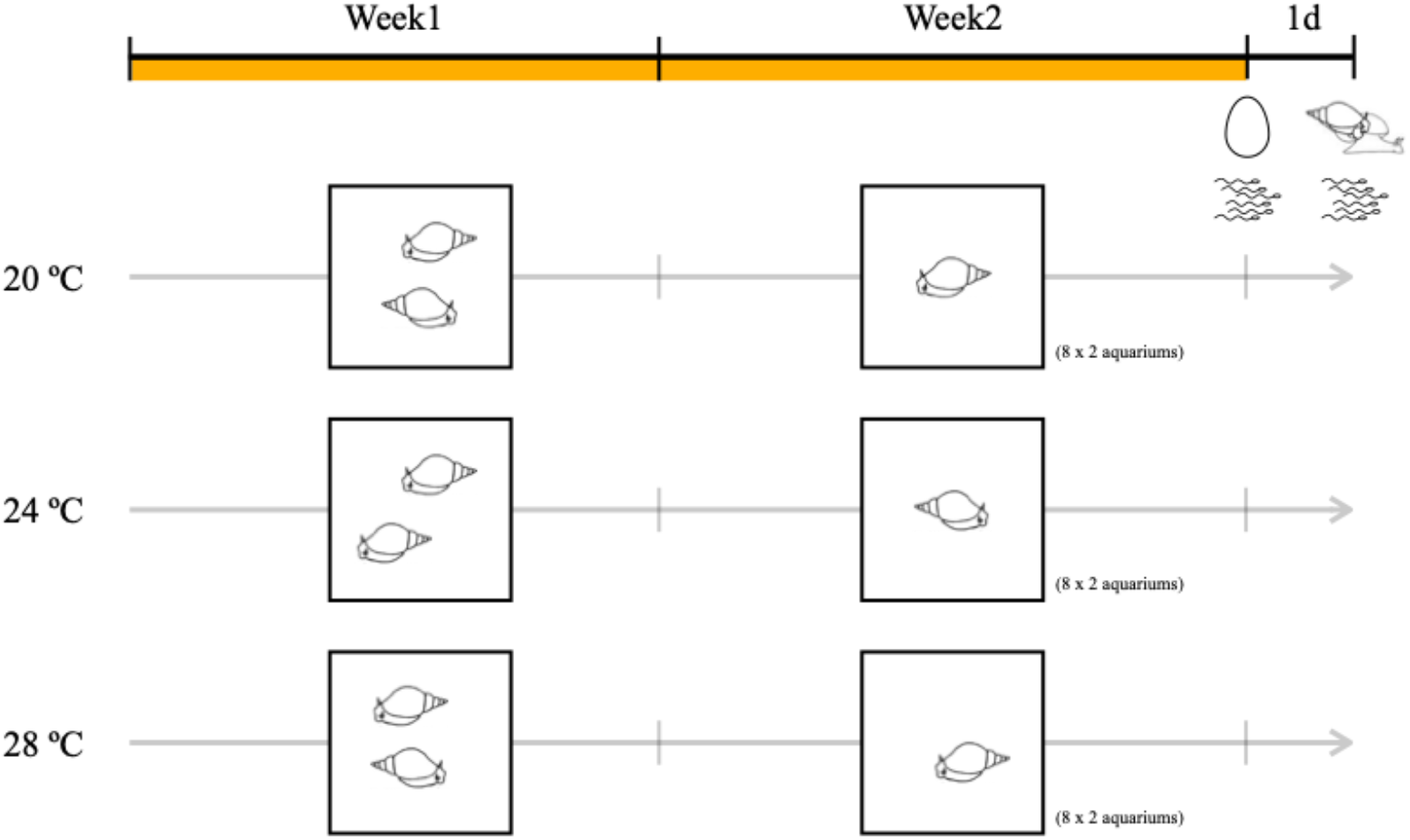
Experimental design. In Week 1, we allowed the snails to mate in pair, so that they used up gametes produced and stored prior to heat treatment. In Week 2, we removed one snail from each container to keep the focal snail isolated. Half of the focal snails was used to measure the production of both types of gametes, as indicated by the pictograms of an egg and sperm, at the end of Week 2. On the following day, we let the other half of the focal snails copulate with the lab snails to measure mating behaviour and sperm transfer (see mating snail and sperm pictogram).

In the first week of heat exposure, we kept two snails together in a container, allowing for depletion of sperm that they had produced and stored in their seminal vesicles before the exposure. We randomly assigned one snail in a pair as focal, by placing a small amount of nail polish on the non-focal individual. Then, at the start of the second week, we removed the non-focal snails from the containers. After exposing the focal snails to the designated temperature for a total of 14 days (see Figure 1), we quantified their sperm and egg productions at the end of Week 2. Furthermore, on Day 15, we provided a mating partner to each focal snail, in order to examine their mating behaviour as well as sperm transfer. In addition, we measured growth of all the focal snails. Based on the sperm and egg production data, we also compared the total investment in reproduction and determined sex allocation across treatments. We describe how we collected these data in the following.

### Growth

Before the start of the temperature treatment, we measured the shell length of focal snails using Vernier callipers. At the end of the exposure, we measured the shell length of focals again to see how much the snails have grown in two weeks.

### Egg production

At the start of Week 2, we provided new containers to each focal individual. At the end of the temperature exposure, we randomly assigned half of the focal snails for measuring egg and sperm production. From the selected focal individuals, we collected all the egg masses laid in Week 2. We scanned the collected egg masses using a flatbed scanner and glass plates with spacers (Canon LiDE 220; see more details in the video instruction in van Iersel et al. 2014). The scanned images of egg masses were used to count the number of egg masses and number of eggs laid using ImageJ (ver. 1.53t, Schneider et al. 2012).

### Sperm production

The focal snails that were used for quantifying egg production, immediately after the treatment, were subsequently used to dissect out their seminal vesicles in order to estimate how many sperm they produced and stored in Week 2. We used the method of sperm counting published, with a few modifications (Loose and Koene 2008). First, we euthanized the snails by slowly injecting ca. 2 ml of 50 mM MgCl_2_ through their foot into the haemocoel using a syringe and needle (30G × ½”). Next, using a coarse forceps, we removed the shell and pinned the soft body onto a dissection plate. Then, using a fine forceps and scissors, we carefully dissected out the whole seminal vesicles and placed these into 800 μl of *Lymnaea* saline solution in a 2 ml tube. Within the solution, we tore apart the duct with a fine forceps and then vortexed for 30 sec. Next, we transferred the duct to a new tube with 400 μl of saline and vortexed it for 30 sec again. We repeated this last step one more time, removed the duct and collected all the solutions into the first tube. After vortexing it for another 30 sec, we took 5 μl of sperm suspension to count the sperm heads, using a Neubauer improved cell counter, and repeated this counting four times for each sample. Lastly, we applied the formula in Loose and Koene (2008) to estimate the number of sperm in the original sperm suspension (depth: 0.1 mm, the number of squares counted: 5, the area of each square: 0.04 mm^2^).

### Mating behaviour

One day after Week 2, we let the remaining set of focal snails copulate with partner snails to measure mating behaviour and sperm transfer. At the end of the temperature treatment, we kept these snails in a flow-through tank at 20 °C for one day. We also isolated partner snails for four days in perforated containers placed in the same flow-through tank. For identification purposes, we put a small amount of nail polish on the shell of partners. Since the focal snails were isolated for eight days, they were fully motivated to copulate as male (Van Duivenboden and Ter Maat 1985; De Boer et al. 1997), and more so than their four-day isolated partners.

On the day of mating observation, we placed one focal and one partner snail together in a container filled with ca. 400 ml of water. The mating observation was conducted for six hours (09:00-15:00) at the control temperature (20 ± 1 °C). According to the series of stereotypical mating behaviours in this species (Koene 2010), we checked all the pairs every 10 min and scored whether they were (i) not in contact, (ii) the focal (or partner) was crawling on the partner’s shell (mounting), (iii) the focal was probing or inseminating. Since insemination usually takes 20-60 min, this sampling interval ensured not missing any copulation.

Based on the mating behaviour data, we counted how many focal individuals mated, and which sex roles they performed first (male or female). We also calculated the duration of mating latency (how long they took to initiate courtship behaviour), that of courtship, and that of insemination. For the analyses of mating behaviour, we only used the cases where focal snails acted as male first, because it is hard to define when they started courtship behaviour if they first mates as females. Moreover, mating as female first has been shown to affects sperm transfer (Nakadera et al. 2014).

### Investment for reproduction and Sex allocation

The change in sperm and egg production in response to temperature further motivated us to examine the change in total reproductive investment as well as sex allocation. To do so, we first made egg and sperm production measurements comparable by standardizing them (mean 0, SD 1) and adding 3 to all the values to make them non-negative. Then, we set that as the total of standardized egg and sperm production as total reproductive investment. For sex allocation, we used the ratio of standardized sperm production divided by total reproductive investment.

### Sperm transfer

Immediately after the focal snail inseminated their partner, we took out the partner and dissected out the vaginal duct which was extensively swollen with ejaculate received (see the method above). We placed the extracted duct in the tube with 400 μl of saline, and tore it apart to release the sperm transferred. We followed the same protocol of sperm counting as above, expect that the total amount of sperm suspension was 1200 μl, not 1600 μl. That is because the number of sperm transferred was expected to be less than that produced.

### Statistics

We carried out all the statistical analyses in R (ver. 4.2.1, R Core Team 2022). Throughout the experiment, we collected the data for growth, egg production, sperm production, sperm transfer and mating behaviour (mating rate, mating role, mating latency, courtship duration, insemination duration). To test if there is any difference between treatments, we applied a statistical model with Treatment (20, 24 and 28 °C) and Run (Run 1 and 2) as fixed factors including interaction. To compare the growth between treatments, we calculated the shell length differences before and after exposure and ran a GLM with Gaussian distribution. To explain the difference we observed, we also ran the same test for the shell length at the start of experiments. To compare the egg production, we used a GLM with Poisson distribution for the number of egg masses, and a GLM with Gaussian distribution for the number of eggs and the number of eggs per egg mass. When there was a significant difference, we applied a Tukey post hoc test. Supplementary, we also compared the number of eggs per egg mass using the same method. To test the sperm production, sperm transfer, total investment for reproduction and sex allocation, we used GLMs with Gaussian distribution and Tukey post hoc tests. For mating rate and mating role, we used GLMs with binomial distribution. Lastly, for the other mating behaviour data, we applied Kruskal-Wallis tests (without including Run as fixed factor), since these variables were not normally distributed.

## Results

Throughout the experiments, two out of 96 snails died in the 28 °C treatment of Run 2. These two snails were excluded from all the analyses.

### Growth

Due to marker loss, the sample sizes for growth data were 29 for 20 °C, 25 for 24°C, and 30 for 28 °C. Within the duration of 2 weeks, we did not detect any difference in growth between treatments (GLM, *F*_2,81_ = 1.88, *P* = 0.159, Fig. 2), although there was a significant difference between Run (GLM, *F*_1,80_ = 11.91, *P* = 0.001, Interaction: *F*_2,78_ = 0.30, *P* = 0.739, Fig. 2). Note that, since we used the same age cohort of snails for both Runs, they had a two-week age difference, and their size of snails in Run 2 was indeed larger at the start of the experiment (GLM, Run: *F*_1,80_ = 24.83, *P* < 0.001, Fig. 2).

**Fig. 2.**
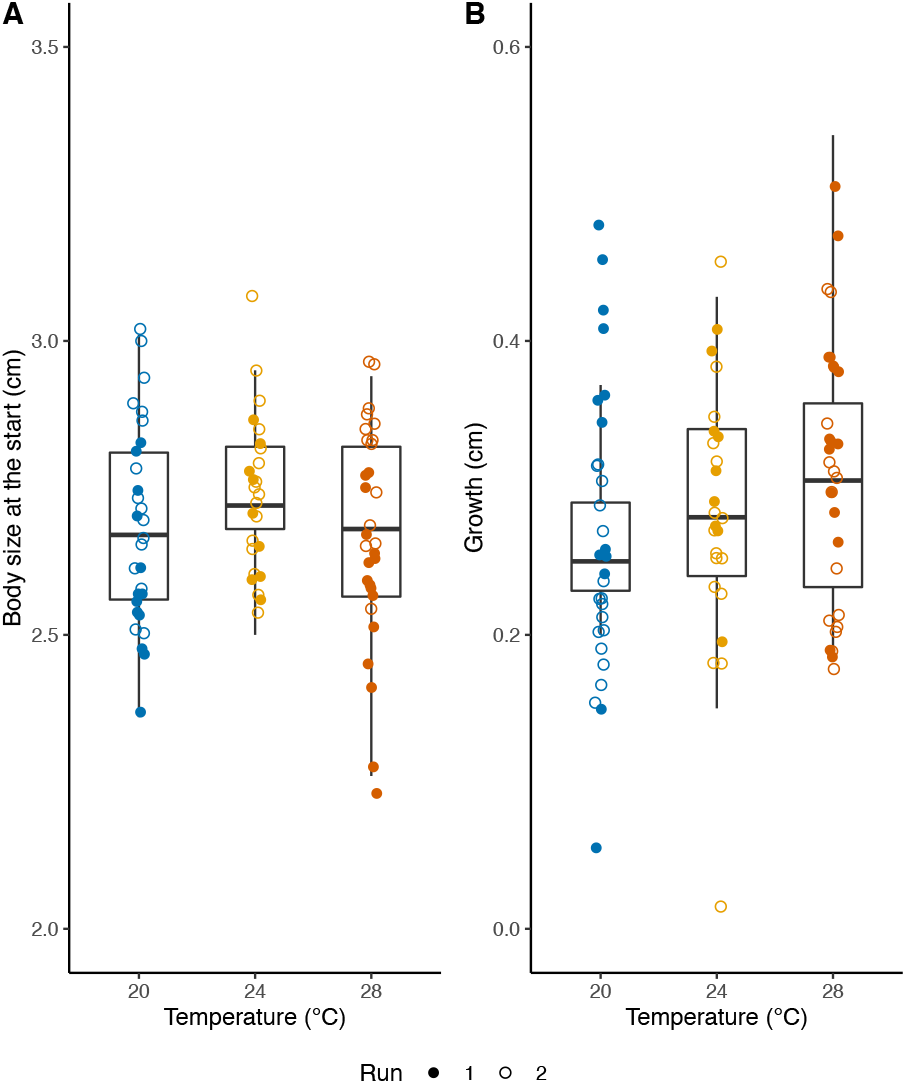
Growth across heat treatments. A. The body size at the start of experiment. The individuals in Run 2 were larger than those in Run 1, since they are 2 weeks older. B. Growth. We plot the difference of shell length at the start and end of experiment as growth. The box plots show median, first and third quartiles and range of data points.

### Egg production

Two snails in the 28 °C treatment did not laid eggs. Therefore, we did not include them to compare the egg production across treatments. There was no difference in the number of egg masses laid across treatments (Fig. 3), but the total number of eggs laid was significantly lower in the snails in 28 °C compared to those in 20 °C (GLM, Treatment: *F*_2,42_ = 7.65, *P* = 0.002, Run: *F*_1, 41_ = 13.36, *P* = 0.001, Interaction: *F*_2,39_ = 2.75, *P* = 0.076, Fig. 3). This was also reflected in the number of eggs per egg mass, which showed the same pattern as the total number of eggs laid (GLM, Treatment: *F*_2,42_ = 6.23, *P* = 0.004, Run: *F*_1,41_ = 10.45, *P* = 0.002, Interaction: *F*_2,39_ = 2.68, *P* = 0.081, Fig. 3). Eventhough not statistically significant, we like to highlight the difference between Runs at 24°C. These two groups of snails had an age difference of two weeks (Fig. 3), and this becomes more prominent in our sex allocation analysis below. Compared to the control (20 °C), egg production was reduced to 40.6 % on average at 28 °C.

**Fig. 3.**
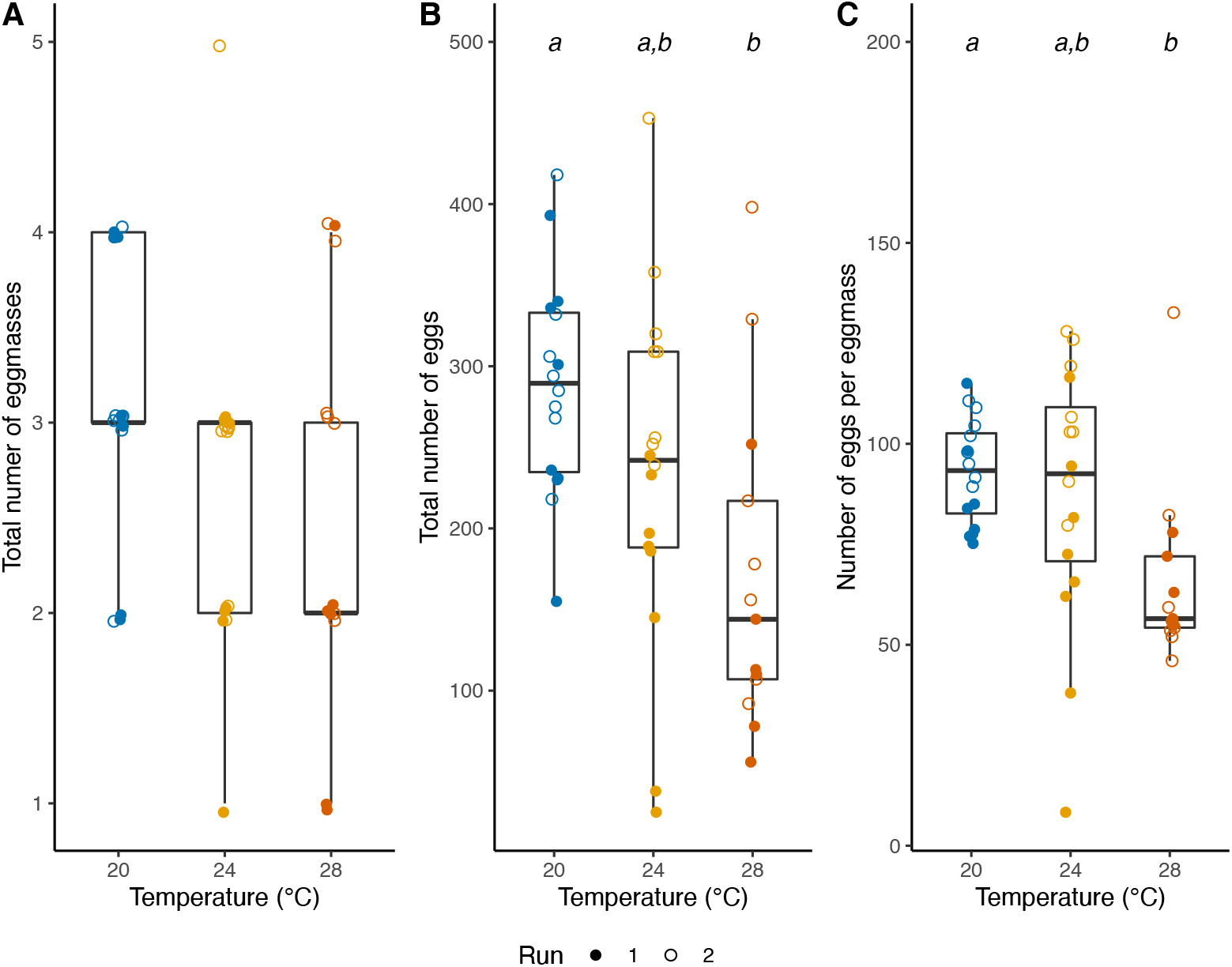
Egg production across heat treatment. A. Total number of egg masses laid in Week 2. B. Total number of eggs laid in Week 2. C. The average number of eggs per egg mass. The letters above box plots indicate the outcome of Tukey post hoc tests. The box plots show median, first and third quartiles and range of data points.

### Sperm production

We detected a significant reduction of sperm production in the snails at 28 °C (GLM, Treatment: *F*_2,44_=82.91, *P* < 0.001, Run: *F*_1,43_ = 0.44, *P* = 0.510, Interaction: *F*_2,41_ = 2.80, *P* = 0.072, Fig. 4a). On average, compared to the control, the snails at 28 °C produced 64.1 % less sperm.

**Fig. 4.**
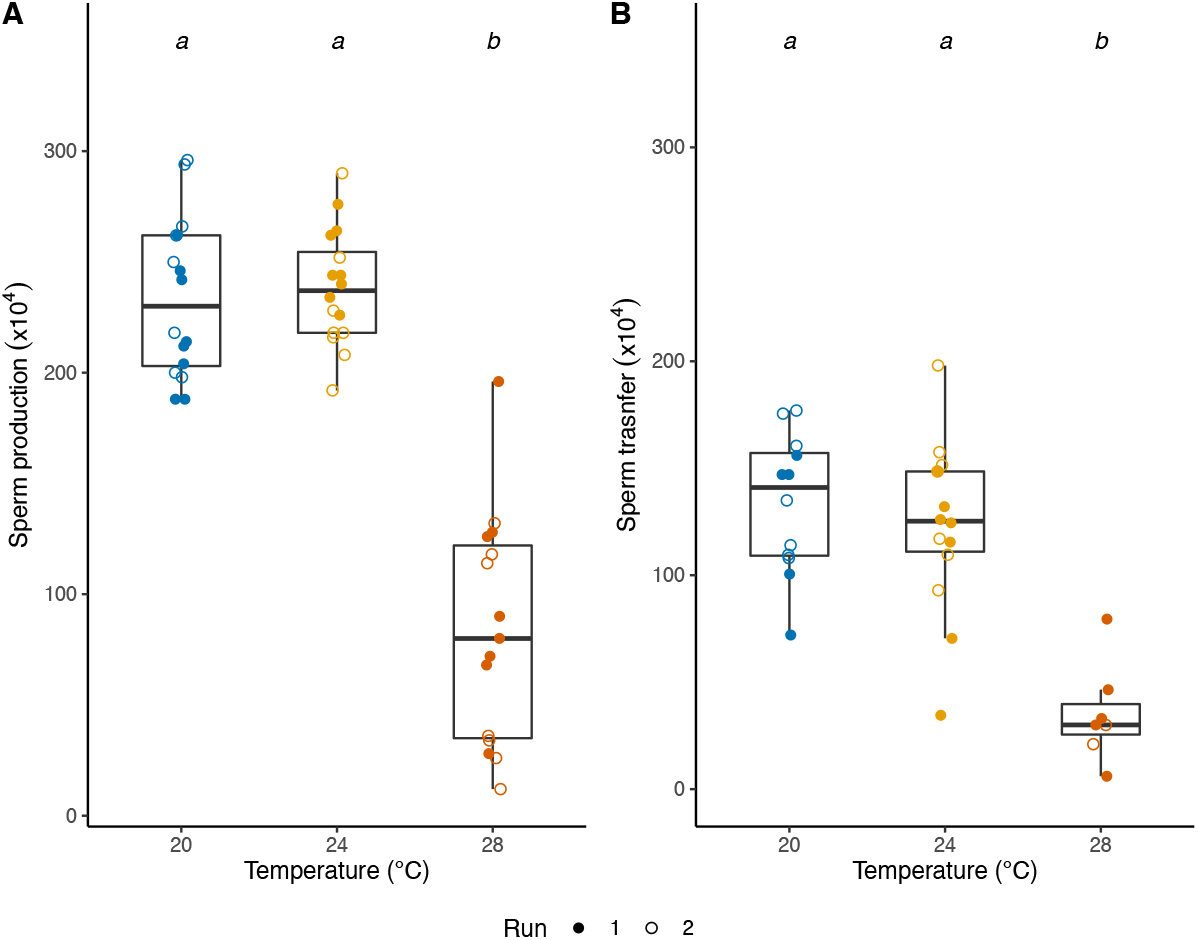
Sperm production across heat treatment. A. The estimated number of sperm produced in Week 2. B. The estimated number of sperm transferred to a mating partner on Week 2 + 1d. The letters above bar plots indicate the outcome of Tukey post hoc tests. The box plots show median, first and third quartiles and range of data points.

### Sperm transfer

Because we can measure sperm transfer only when the focal snails had copulated, the sample size is smaller and varying (Run1: N = 17, Run2: N = 16, 20 °C: N = 12, 24°C: N = 14, 28 °C: N = 7). The data show a very similar pattern to the sperm production results: the snails at 28 °C transferred significantly less sperm to their mating partner (Treatment: *F*_2,30_=20.88, *P* < 0.001, Run: *F*_1,29_ = 2.08, *P* = 0.160, Interaction: *F*_2,27_ = 0.91, *P* = 0.413, Fig. 4b). On average, compared to the control, the snails at 28 °C transferred 73.7 % less sperm.

### Mating behaviour

The mating rate did not differ between treatments, when they mated with non-heat-treated, control partners (Treatment: *χ*^*2*^_2_ = 1.96, *P* = 0.098, Fig. 5). Their mating roles also did not differ significantly between treatments (Treatment: *χ*^*2*^_2_ = 3.31, *P* = 0.354, Fig. 5). For the remaining mating behaviour data, we had to exclude one sample (Run 2, 28 °C) that we prematurely interrupted for sperm counting; the focal snail had not transferred an ejaculate yet. Nonetheless, we did not find any significant difference in mating latency (Kruskal-Wallis test, *χ*^*2*^_2_ = 1.16, *P* = 0.559, Fig. 6), courtship duration (Kruskal-Wallis test, *χ*^*2*^_2_ = 1.85, *P* = 0.397, Fig. 6) and insemination duration (Kruskal-Wallis test, *χ*^*2*^_2_ = 1.73, *P* = 0.420, Fig. 6).

**Fig. 5.**
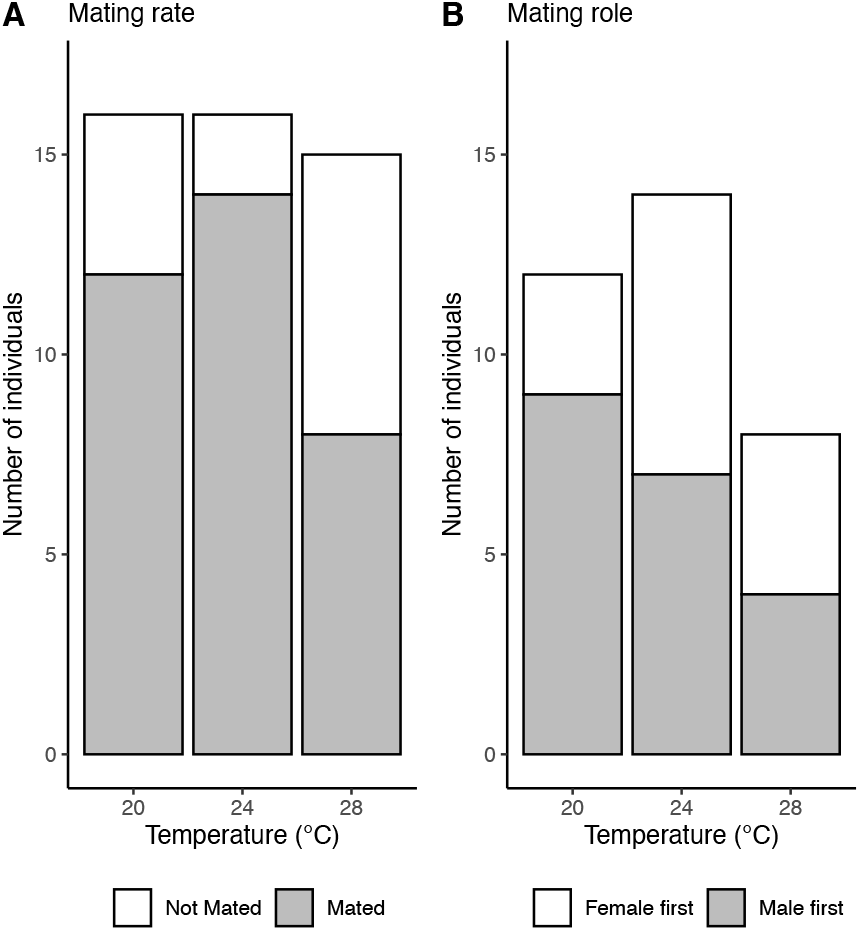
A. Mating rate after heat treatment. We counted how many focal individuals mated with control, non-heat-treated snails. B. Mating role after heat treatment. We counted how many focal snails acted as male or female in their first mating with control snails.

**Fig. 6.**
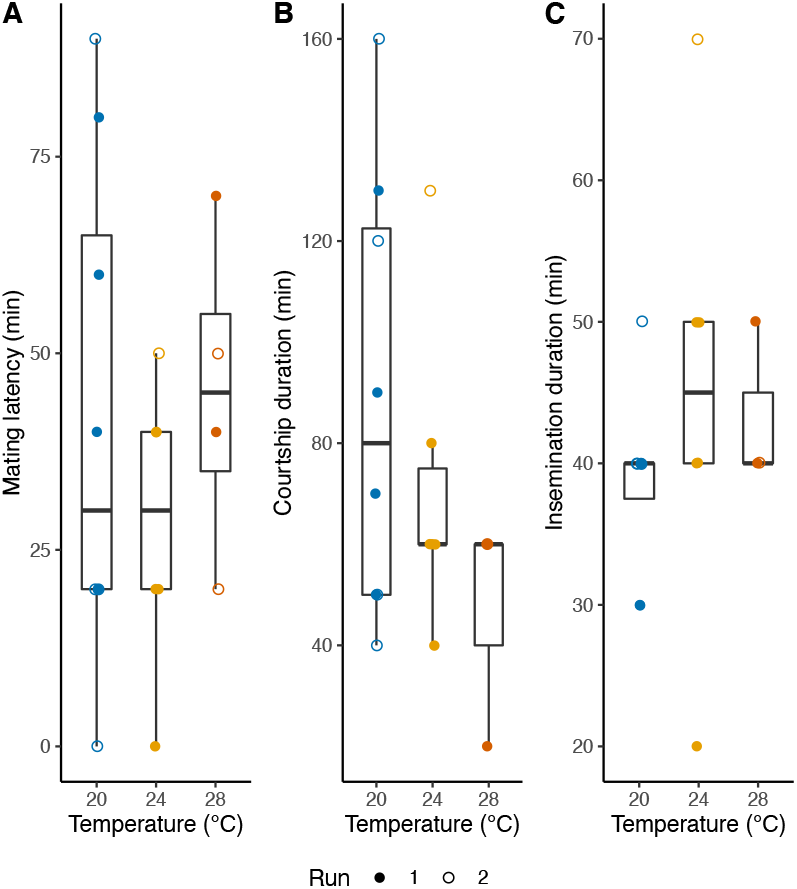
Mating behaviours of heat-treatment snails mating with control partners. We measured the male mating behaviour of focal snails when they acted as male first. A. Mating latency. We plotted the duration from the start of mating trial until a focal snail initiated male courtship behaviour. B. Courtship duration. We measured the duration of courtship from mounting to the end of probing. C. Insemination duration. The y-axis indicates the duration of ejaculate transfer. The box plots show median, first and third quartiles and range of data points.

### Investment for reproduction and sex allocation

There was a significant reduction in total reproductive investment in the 28 °C treatment (Treatment: *F*_2,42_ = 44.51, *P* < 0.001, Run: *F*_1,41_ = 8.61, *P* = 0.006, Interaction: *F*_2,39_ = 0.66, *P* = 0.521, Fig. 7). Sex allocation was also different across treatments and as well as Run and interaction (Treatment: *F*_2,42_ = 12.82, *P* < 0.001, Run: *F*_1,41_ = 13.54, *P* = 0.001, Interaction: *F*_2,39_ = 4.35, *P* = 0.027, Fig. 7). The male allocation in the snails exposed to 28 °C was significantly reduced, compared to the control treatment. The interaction was most likely due to the variation between Runs in the 24 °C treatment (Fig. 7), suggesting that this is an effect of age, as also shown in egg production at 24 °C.

**Fig. 7.**
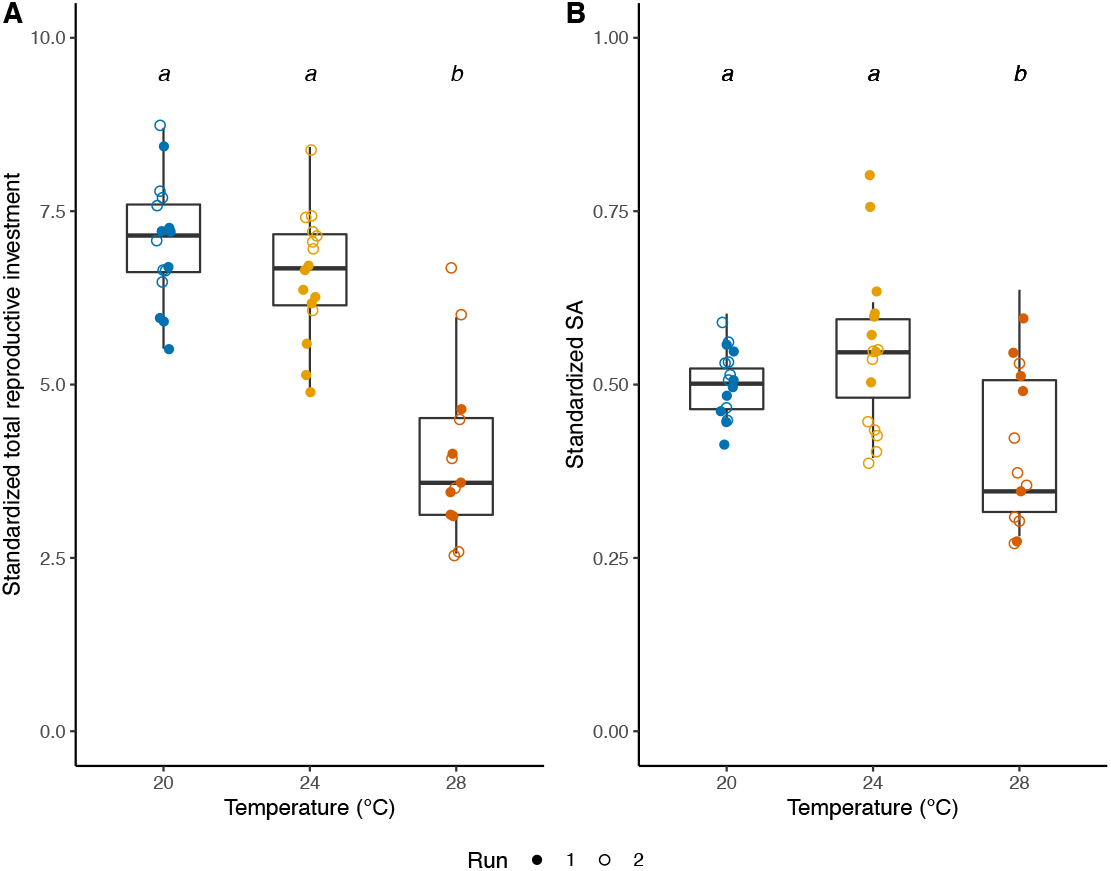
Investment for reproduction and sex allocation. The letters above bar plots indicate the outcome of Tukey post hoc tests. The box plots show median, first and third quartiles and range of data points.

## Discussion

We found that exposing snails to sub-lethal temperature significantly reduced both egg and sperm productions, and that their male function was more vulnerable than their female function. Also, we found that, despite of their reduced sperm production and transfer, their male mating motivation was not affected. Lastly, we did not observe any effect of heat on growth. In the following, we will discuss these findings and place them in a broader ecological and evolutionary perspective.

The control group invested equally in male and female reproduction, which is in accordance with previous studies looking at sex allocation in this species (Koene et al. 2006; Hoffer et al. 2010). We found that the total investment in reproduction especially declined in the snails exposed to 28 °C. This pattern supports the consensus that reproduction is vulnerable to sublethal heat exposure (Walsh et al. 2019). Our results also indicate that snails exposed to 28 °C allocated proportionally more to their female function, compared to the control snails (Fig. 7). The shifted sex allocation and diminished sperm production and transfer leads us to conclude that the male function is more vulnerable to heat stress in this species. We designed the experiment with the aim to measure the consequence of heat on spermatogenesis, even though this species continuously produces sperm after maturation. sPrevious work has shown that one week of isolation suffices to fully replenish the components of ejaculate in this species, increasing their male mating motivation (Van Duivenboden and Ter Maat 1985; De Boer et al. 1997). Thus, we are confident that snails in 28 °C treatment did not fully replenish their sperm reserves. Clearly, it remains to be determined which spermatogenesis stage is affected and whether the produced sperm are viable. In addition, the observed interaction with heat treatment and experimental runs in sex allocation implies that the temperature vulnerability may be age specific. That is, even though they are only two weeks apart, the younger snails might be more vulnerable.

We found that sperm production was reduced under heat and this reduction strongly influenced the sperm transfer of *L. stagnalis* (Fig. 4), but not their mating motivation and behaviour (Fig. 5, 6). Since this species transfers sperm at the end of insemination (Weggelaar et al. 2019), it is expected to have the disassociation between the number of sperm transferred and insemination duration. Also, we like to emphasize that the snails at 20 °C and 24 °C transferred approximately 50% of sperm stored, and the snails at 28 °C used almost all sperm they had (Fig. 4). This pattern implies that the snails know how many sperm they have in seminal vesicles; for a different species it has been shown that sperm release can be controlled by the donor (Geoffroy et al. 2005). However, their male mating motivation was not affected as drastically as one would imagine from their reduced sperm production (Fig. 4).

Crucially, such unaffected male mating motivation in *L. stagnalis* was also observed in a different context. When snails receive seminal fluid proteins, they significantly reduce the number of sperm transferred in a subsequent mating (Nakadera et al. 2014), although their male mating motivation stays unchanged (Nakadera et al. 2015). The observed mismatch of reduced sperm production and unchanged male mating motivation in this study is particularly concerning for field populations, since the heated snails would waste energy by transferring a few sperm, while facing the higher risk of infection due to compromised immune defence (e.g., Seppälä and Jokela 2011; Leicht et al. 2013). We like to stress that we chose to let the snails have a day at 20 °C before mating, to test the effect of reduced sperm production on mating behaviour. This ‘rest’ day might have affected their male mating motivation and behaviour. For future research, it would be interesting to directly examine how heat affects mating behaviour.

We did not find an effect of heat on growth in our experimental design. However, it is likely that, if we had kept the snails under heat treatment for a longer period, we would have detected such a difference (e.g., Leicht et al. 2013; Hoefnagel and Verberk 2017; Salo et al. 2019). The trend that the snails at 28 °C show might be seen as a confirmation for this previously reported response, and is not surprising given their increased metabolic rate due to the temperature increase. Even though there is a positive correlation between body size and female fecundity in this species (Koene et al. 2007), we observed a significant reduction in egg production under heat (Fig. 3B, also see Leicht et al. 2013; Salo et al. 2019). This reduction probably occurs because the heated snails deposited less eggs in an egg mass, rather than changing their egg laying frequency (Fig. 3C). Similar to sperm production, we did not measure the quality of eggs produced under heat, and do not know which stage of egg production was affected by heat. Since *L. stagnalis* can store and use sperm from mating partners for ca. three months (Cain 1956; Nakadera et al. 2014b), the snails should have had plenty of sperm to fertilize during the treatment, although we cannot exclude the possibility that the stored sperm got deteriorated under heat.

This study demonstrated that examining hermaphrodites provides vital insights on the sex differences under heat, which is not accessible in separate-sexed species. As commonly expected or assumed in diverse species (Sage et al. 2015; Walsh et al. 2019a,b; Iossa 2019), the male function of *L. stagnalis* is more sensitive to elevated temperature, which means that the proxy mechanism of those responses can be shared across wide range of species. As many hermaphrodites, *L. stagnalis* produces sperm and eggs in a same organ called ovotestis (Davison 2006; Koene et al., 2006).

Although the ovotestis is a particularly interesting target organ, its current understanding in gastropods is relatively limited, in terms of gene expression, distribution of oo- and spermatogenesis sites, or the fate determination of germ cells. This study paved the path to investigate the proximate mechanisms of reduced male and female fertility under heat in a hermaphroditic species and to predict the implications in natural populations. Moreover, with temperature projected to increase in future, we hope this study motivates further studies investigating the impact of heat exposure in wide range of hermaphrodites.

## Acknowledgement

We thank Omer Ballaoui and Esther D. Hoekman for setting the aquariums up and maintaining the snail culture in the lab, and Angus Davison and Simultaneous Hermaphroditic Organism Workshop (SHOW) community for inspirations and encouragements (including Lukas Schärer, Cynthia Norton, Janet Leonard, Steven A. Ramm, Maria Christina Lorenzi, Alexandra Staikou, Chiara Benvenuto).

## Author contributions

Conceptualization: Y.N., J.M.K., V.Z.; Methodology: Y.N., J.M.K., V.Z.; Formal analyses: Y.N., S.v.D.; Investigation: S.v.D., Y.N.; Data curation S.v.D.; Writing – original draft: Y.N.; Writing – review and editing: Y.N., J.M.K., V.Z.; Supervision: Y.N., J.M.K., V.Z.; Project administration: J.M.K.

## Reference

Blanckenhorn, W. U., Gautier, R., Nick, M., Puniamoorthy, N., & Schäfer, M. A. (2014). Stage- and sex-specific heat tolerance in the yellow dung fly Scathophaga stercoraria. Journal of Thermal Biology, 46, 1–9.

Cain, G. L. (1956). Studies on cross-fertilization and self-fertilization in Lymnaea stagnalis Appressa Say. The Biological Bulletin, 111, 45–52.

Charnov, E. L. (1979). Simultaneous hermaphroditism and sexual selection. Proceedings of the National Academy of Sciences of the United States of America, 76, 2480–2484.

Davison, A. (2006). The ovotestis: An underdeveloped organ of evolution. BioEssays, 28, 642–650.

De Boer, P. A. C. M., Jansen, R. F., Koene, J. M., & Ter Maat, A. (1997). Nervous control of male sexual drive in the hermaphroditic snail Lymnaea stagnalis. Journal of Experimental Biology, 951, 941–951.

De Jong-Brink, M., Jager, J. C., Bolwerk, E. L. M., & Jong, J. T. L. (1985). Statistical analysis of frequencies and proportions of cytologically classified spermatogenesis cells in the hermaphroditic snail Lymnaea stagnalis. 1. A study of diurnal variations. International journal of invertebrate Reproduction and Development, 8, 149–159.

Doums, C., & Jarne, P. (1996). The evolution of phally polymorphism in Bulinus truncatus (Gastropoda, Planorbidae): the cost of male function analysed through life-history traits and sex allocation. Oecologia, 106(4), 464–469.

Doums, C., Perdieu, M.-A., & Jarne, P. (1998). Resource allocation and stressful conditions in the aphallic snail Bulinus truncatus. Ecology, 79(2), 720–733.

Geoffoy, E., Hutcheson, R., & Chase, R. (2005). Nervous control of ovulation and ejaculation in Helix aspersa. Journal of molluscan studies, 71(4), 393–399.

Hoefnagel, K. N., & Verberk, W. C. E. P. (2017). Long-term and acute effects of temperature and oxygen on metabolism, food intake, growth and heat tolerance in a freshwater gastropod. J Therm Biol, 68(Pt A), 27–38.

Hoffer, J. N. A., Ellers, J., & Koene, J. M. (2010). Costs of receipt and donation of ejaculates in a simultaneous hermaphrodite. BMC evolutionary biology, 10, 393.

Hurley, L. L., McDiarmid, C. S., Friesen, C. R., Griffith, S. C., & Rowe, M. (2018). Experimental heatwaves negatively impact sperm quality in the zebra finch. Proceedings of the Royal Society B: Biological Sciences, 285.

Iossa, G. (2019). Sex-Specific Differences in Thermal Fertility Limits. Trends in Ecology and Evolution, 34, 490–492.

Koene, J. M. (2010). Neuro-endocrine control of reproduction in hermaphroditic freshwater snails: mechanisms and evolution. Frontiers in behavioral neuroscience, 4, 167.

Koene, J. M., & Ter Maat, A. (2005). Sex role alternation in the simultaneously hermaphroditic pond snail Lymnaea stagnalis is determined by the availability of seminal fluid. Animal Behaviour, 69, 845–850.

Koene, J. M., Montagne-Wajer, K., & Ter Maat, A. (2006). Effects of frequent mating on sex allocation in the simultaneously hermaphroditic great pond snail (Lymnaea stagnalis). Behavioral Ecology and Sociobiology, 60, 332–338.

Lee Teskey, M., Lukowiak, K. S. K., Riaz, H., Dalesman, S., & Lukowiak, K. S. K. (2012). Research article: What’s hot: The enhancing effects of thermal stress on long-term memory formation in Lymnaea stagnalis. Journal of Experimental Biology, 215, 4322–4329.

Leicht, K., Jokela, J., & Seppälä, O. (2013). An experimental heat wave changes immune defense and life history traits in a freshwater snail. Ecology and Evolution, 3, 4861–4871.

Leicht, K., Seppälä, K., & Seppälä, O. (2017). Potential for adaptation to climate change: Family-level variation in fitness-related traits and their responses to heat waves in a snail population. BMC Evolutionary Biology, 17, 1–10.

Leicht, K., & Seppälä, O. (2019). Direct and transgenerational effects of an experimental heatwave on early life stages in a freshwater snail. Freshwater Biology, 64, 2131–2140.

Leicht, K., Jokela, J., & Seppälä, O. (2019). Inbreeding does not alter the response to an experimental heat wave in a freshwater snail. PLoS ONE, 14, 1–12.

Loose, M. J., & Koene, J. M. (2008). Sperm transfer is affected by mating history in the simultaneously hermaphroditic snail Lymnaea stagnalis. Invertebrate Biology, 127, 162–167.

Moussaoui, R., Verdel, K., Benbellil-Tafoughalt, S., & Koene, J. M. (2018). Female behaviour prior to additional sperm receipt in the hermaphroditic pond snail Lymnaea stagnalis. Invertebrate Reproduction and Development, 62, 82–91.

Nakadera, Y., Blom, C., & Koene, J. M. (2014). Duration of sperm storage in the simultaneous hermaphrodite Lymnaea stagnalis. Journal of Molluscan Studies, 80, 1–7.

Nakadera, Y., Swart, E. M., Hoffer, J. N. A., den Boon, O., Ellers, J., Koene, J. M. (2014). Receipt of seminal fluid proteins causes reduction of male investment in a simultaneous hermaphrodite. Current biolog : CB, 24, 859–862.

Nakadera, Y., Swart, E. M., Maas, J. P. A., Montagne-Wajer, K., Ter Maat, A., & Koene, J. M. (2015). Effects of age, size, and mating history on sex role decision of a simultaneous hermaphrodite. Behavioral Ecology, 26, 232–241.

Nakadera, Y., Thornton Smith, A., Daupagne, L., Coutellec, M., Koene, J. M., & Ramm, S. A. (2020). Divergence of seminal fluid gene expression and function among natural snail populations. Journal of Evolutionary Biology, 33, 1440–1451.

Oshino, T., Abiko, M., Saito, R., Ichiishi, E., Endo, M., Kawagishi-Kobayashi, M. et al. (2007). Premature progression of anther early developmental programs accompanied by comprehensive alterations in transcription during high-temperature injury in barley plants. Mol Genet Genomics, 278(1), 31–42.

Park, C. B., Kim, Y. J., & Soyano, K. (2017). Effects of increasing temperature due to aquatic climate change on the self-fertility and the sexual development of the hermaphrodite fish, Kryptolebias marmoratus. Environmental Science and Pollution Research, 24, 1484–1494.

Parratt, S. R., Walsh, B. S., Metelmann, S., White, N., Manser, A., Bretman, A. J. et al. (2021). Temperatures that sterilize males better match global species distributions than lethal temperatures. Nature Climate Change, 11(6), 481–484.

Paxton, C. W., Baria, M. V., Weis, V. M., & Harii, S. (2016). Effect of elevated temperature on fecundity and reproductive timing in the coral Acropora digitifera. Zygote, 24(4), 511–516.

Prasad, A., Croydon-Sugarman, M. J., Murray, R. L., & Cutter, A. D. (2011). Temperature-dependent fecundity associates with latitude in Caenorhabditis briggsae. Evolution, 65(1), 52–63.

Pélissié, B., Jarne, P., & David, P. (2012). Sexual selection without sexual dimorphism: Bateman gradients in a simultaneous hermaphrodite. Evolution 66, 66–81.

Sage, T. L., Bagha, S., Lundsgaard-Nielsen, V., Branch, H. A., Sultmanis, S., & Sage, R. F. (2015). The effect of high temperature stress on male and female reproduction in plants. Field Crops Research, 182, 30–42.

Sales, K., Vasudeva, R., Dickinson, M. E., Godwin, J. L., Lumley, A. J., Michalczyk, Ł. et al. (2018). Experimental heatwaves compromise sperm function and cause transgenerational damage in a model insect. Nature Communications, 9, 4771.

Salo, T., Stamm, C., Burdon, F. J., Räsänen, K., & Seppälä, O. (2017). Resilience to heat waves in the aquatic snail Lymnaea stagnalis: Additive and interactive effects with micropollutants. Freshwater Biology, 62, 1831–1846.

Salo, T., Kropf, T., Burdon, F. J., & Seppälä, O. (2019). Diurnal variation around an optimum and near-critically high temperature does not alter the performance of an ectothermic aquatic grazer. Ecology and Evolution, 9, 11695–11706.

Schneider, C. A., Rasband, W. S., & Eliceiri, K. W. (2012). NIH Image to ImageJ: 25 years of image analysis. Nature methods, 9(7), 671–675.

Schrag, S. J., & Read, A. F. (1992). Temperature determination of male outcrossing ability in a simultaneous hermaphrodite. Evolution, 46(6), 1698–1707.

Schärer, L. (2009). Tests of sex allocation theory in simultaneously hermaphroditic animals. Evolution, 63, 1377–1405.

Seppälä, O., & Jokela, J. (2011). Immune defence under extreme ambient temperature. Biology Letters, 7, 119–122.

Sidorov, A. V. (2002). Effect of temperature on synaptic transmission between identified neurones of the mollusc Lymnaea stagnalis. Neuroscience letters, 333(1), 1–4.

Sidorov, A. V. (2012). Temperature dependence of monoamine-induced pulmonary respiration in the mollusc Lymnaea stagnalis. Journal of Evolutionary Biochemistry and Physiology, 48(3), 287–294.

Steen, W. J. V. D., Hoven, N. P. V. D., & Jager, J. C. (1968). A method for breeding and studying freshwater snails under continuous water change, with some remarks on growth and reproduction in Lymnaea stagnalis (L.). Netherlands Journal of Zoology, 19, 131–139.

Stürup, M., Baer-Imhoof, B., Nash, D. R., Boomsma, J. J., & Baer, B. (2013). When every sperm counts: Factors affecting male fertility in the honeybee Apis mellifera. Behavioral Ecology, 24, 1192–1198.

Sunada, H., Riaz, H., De Freitas, E., Lukowiak, K. K., Swinton, C., Swinton, E. et al. (2016). Heat stress enhances LTM formation in Lymnaea: Role of HSPs and DNA methylation. Journal of Experimental Biology, 219, 1337–1345.

Van Duivenboden, Y. A., & Ter Maat, A. (1985). Masculinity and receptivity in the hermaphrodite pond snail, Lymnaea stagnalis. Animal Behaviour, 33, 885–891.

Van Iersel, S., Swart, E. M., Nakadera, Y., van Straalen, N. M., & Koene, J. M. (2014). Effect of male accessory gland products on egg laying in gastropod molluscs. Journal of visualized experiments: JoVE, 88, e51698.

Vaughn, C. M. (1953). Effects of Temperature on Hatching and Growth of Lymnaea stagnails appressa Say. The American Midland Naturalist, 49, No. 1(Jan., 1953), 214–228.

Walsh, B. S., Parratt, S. R., Hoffmann, A. A., Atkinson, D., Snook, R. R., Bretman, A. et al. (2019). The Impact of Climate Change on Fertility. Trends in Ecology and Evolution, 34, 249–259.

Walsh, B. S., Parratt, S. R., Atkinson, D., Snook, R. R., Bretman, A., & Price, T. A. R. (2019). Thermal fertility limits and sex: an integrated approach. Trends in Ecology and Evolution.

Weggelaar, T. A., Commandeur, D., & Koene, J. M. (2019). Increased copulation duration does not necessarily reflect a proportional increase in the number of transferred spermatozoa. Animal Biology, 69, 95–115.

Yogev, L., Kleiman, S., Shabtai, E., Botchan, A., Gamzu, R., Paz, G. et al. (2004). Seasonal variations in pre- and post-thaw donor sperm quality. Hum Reprod, 19(4), 880–885.

Zhang, P., Blonk, B. A., van den Berg, R. F., & Bakker, E. S. (2018). The effect of temperature on herbivory by the omnivorous ectotherm snail Lymnaea stagnalis. Hydrobiologia, 812(1), 147–155.

Zizzari, Z. V., & Ellers, J. (2011). Effects of exposure to short-term heat stress on male reproductive fitness in a soil arthropod. Journal of Insect Physiology, 57, 421–426.

Zizzari Z.V., Ellers J. 2014. Rapid shift in thermal resistance between generations through maternal heat exposure. Oikos, 123:1365–1370.

